# Spatiotemporal analysis of *Plasmodium berghei* sporozoite maturation and infectivity in *Anopheles* vector

**DOI:** 10.1101/2025.07.21.665974

**Authors:** Lisa H Verzier, Jean-Michel Thiberge, Eduardo Aliprandini, Vanessa Lagal, Pauline Formaglio, Olivier Silvie, Rogerio Amino

## Abstract

*Plasmodium* sporozoites must undergo tightly regulated developmental transitions to become infectious and be successfully transmitted from the mosquito vector to a mammalian host. While transcriptomic studies have revealed stage-specific changes across sporozoite populations, the functional consequences of these transitions remain unclear. Here, using *P. berghei*, we characterised over time the infectivity of sporozoite forms collected from the midgut, haemolymph, salivary glands and saliva. We show that salivary gland invasion is required but not sufficient for sporozoite optimal infectivity, with the acquisition of hepatocyte cell traversal and invasion progressively increasing until a plateau from 18 days post-infection onwards. Using a stage-specific fluorescent reporter to track maturation, we found that fluorescence increased over time and correlated with infectivity within each compartment, yet maximal infectivity was reached only in salivary gland sporozoites. Notably, our data suggest that salivated sporozoites—the natural transmission form—exhibit enhanced infectivity relative to gland-resident forms both *in vitro* and *in vivo* early after salivary gland invasion. This difference decreases following optimal maturation inside the glands over time. These observations show a crescent gradient of sporozoite maturation and infectiveness from the midgut to the saliva, in which pre-salivary gland forms retain limited infectivity but do not reach the competence of mature salivary gland-associated parasites.

## Introduction

*Plasmodium* parasites have evolved to transition between two radically different environments as they cycle between vertebrate and invertebrate hosts. Transiting from a warm-blooded mammal to the mosquito and vice-versa, they must adapt to hosts with distinct metabolic demands and immune landscapes. While transmission to the mosquito triggers the rapid differentiation of gametocytes into gametes, return to the mammalian host depends entirely on a single, multi-invasive specialised form, the sporozoite (SPZ).

SPZ are formed within oocysts nestled under the basal lamina of the mosquito midgut. Upon maturation, they egress into the haemolymph—a process thought to involve protease secretion and activation of gliding motility (1–3). They are then passively carried through the haemolymph until they encounter the salivary glands, where they specifically recognise and invade acinar cells (4–7). Salivary gland invasion is a complex, multi-step process. SPZ must breach the basal lamina, traverse acinar cells via transient vacuoles or membrane rupture, and ultimately enter the secretory cavities (8–11). While most parasites accumulate in a quiescent state within the secretory cavities, a minority reach the salivary ducts, becoming candidates for injection into the mammalian dermis during the next bloodmeal (12–14). These anatomical distinctions have led to the operational classification of SPZ into oocyst or midgut (MG), haemolymph (HL), and salivary gland (SG) forms.

The SPZ infectivity—defined as the ability of a pathogen to enter, survive, and replicate within a host (15)—entails the parasite’s capacity to navigate host tissues, traverse cells, and invade and develop intracellularly inside a targeted cell. Following deposition by an infected *Anopheles* mosquito, typically in the dermis, SPZ display rapid motility (>1 µm/s) that enables them to leave the injection site and locate blood vessels (16–18). Once in circulation, SPZ are transported to the liver where they cross the sinusoidal barrier via endothelial or Kupffer cell traversal, or via a cell traversal independent mechanism, ultimately reaching the hepatic parenchyma (4,19–22). After traversal of hepatocytes, yet-unknown factors trigger invasion and subsequent differentiation into the exoerythrocytic form (EEF) (23–28). Motility, cell traversal, and hepatocyte invasion are therefore the principal cellular determinants of SPZ infectivity.

The question of when and how sporozoites acquire infectivity has been the subject of long-standing investigation since the 1930s (29–31). Early experimental work had already shown that sporozoites can infect the vertebrate host before direct contact with the mosquito salivary glands (32). Vanderberg later showed that SG SPZ were more infectious than HL and MG SPZ and proposed that infectivity acquisition was driven primarily by time spent in the mosquito rather than by tissue-specific cues (33). This interpretation was subsequently challenged by evidence that MG SPZ could colonise SG in naïve mosquitoes, whereas SG SPZ could not re-invade, pointing instead to tissue-specific developmental programming (34). More recently, Sato et al. showed that HL SPZ are already infectious and can elicit protective immunity, yet remain functionally impaired relative to SG SPZ. Corroborating the compartment-specific maturation model, transcriptomics comparing MG and SG SPZ identified genes “up-regulated in infective sporozoites” (UIS) and “up-regulated in oocyst sporozoites” (UOS) (35–38). Genes included in the former list include UIS3 and UIS4 which are essential in liver stage development, while TREP (UOS3), CRMP2 and CRMP4 are essential for egress from oocyst and SG invasion (36,39–42). Additional transcriptomic and proteomic profiling of SPZ populations has revealed significant differences across forms, including the discovery of translational repression mechanisms during SPZ maturation (14,28,37,38,43). Critical liver-stage proteins such as UIS3 and UIS4 were identified through such comparative approaches. More recently, single-cell RNA-sequencing studies have provided fine-grained resolution of SPZ transcriptional heterogeneity across forms (44–46). Notably, Real and colleagues reported a major transcriptional shift between *P. falciparum* oocyst and HL SPZ, while Bogale and co-workers observed a larger shift between HL and SG SPZ in *P. berghei*. Whether these discrepancies reflect species differences, collection techniques, or biological variation remains unresolved.

Despite these molecular insights, how transcriptomic and proteomic changes translate into functional infectivity remains poorly defined. A detailed understanding of how sporozoites acquire infectivity is not only fundamental to malaria transmission biology but also critical for the rational development of axenic sporozoites, which has yet to replicate this maturation process outside the mosquito vector (47,48). Moreover, most datasets represent single timepoints and do not capture the temporal dimension of SPZ maturation. The role of salivation itself—the physiological route of transmission—also remains understudied. To address these gaps, we examined the infectivity of *P.berghei* SPZ collected across different compartments and timepoints using complementary *in vitro* and *in vivo* assays. Our findings reveal that both tissue and duration of residence influence SPZ competence and identify a transcription marker as a proxy for maturation. Additionally, we uncover that salivated SPZ—the natural form transmitted to mammals—display enhanced infectivity relative to SG SPZ, particularly during early mosquito infection, highlighting the biological relevance of this neglected population.

## Results

### Sporozoites acquire optimal infectivity following salivary gland invasion

*Plasmodium* SPZ are formed within oocysts located between the basal lamina surrounding the mosquito MG and the epithelial layer. Upon maturation, they egress into the HL and subsequently invade the SG, where they reside quiescent, possibly for several days, prior to transmission to a mammalian host during a blood meal (Fig. 1a). To correlate transcriptomic insights with phenotypic evidence, we systematically characterised the infectivity of these SPZ populations by quantifying their capacity for hepatocyte invasion and cell traversal, measured respectively by scoring the percentage of fluorescent SPZ inside HepG2 hepatoma cells and by the percentage of wounded and resealed HepG2 cells after 2 hours of incubation with SPZ collected over the course of mosquito infection (12-28 days post-infection (pi)).

**Figure 1.**
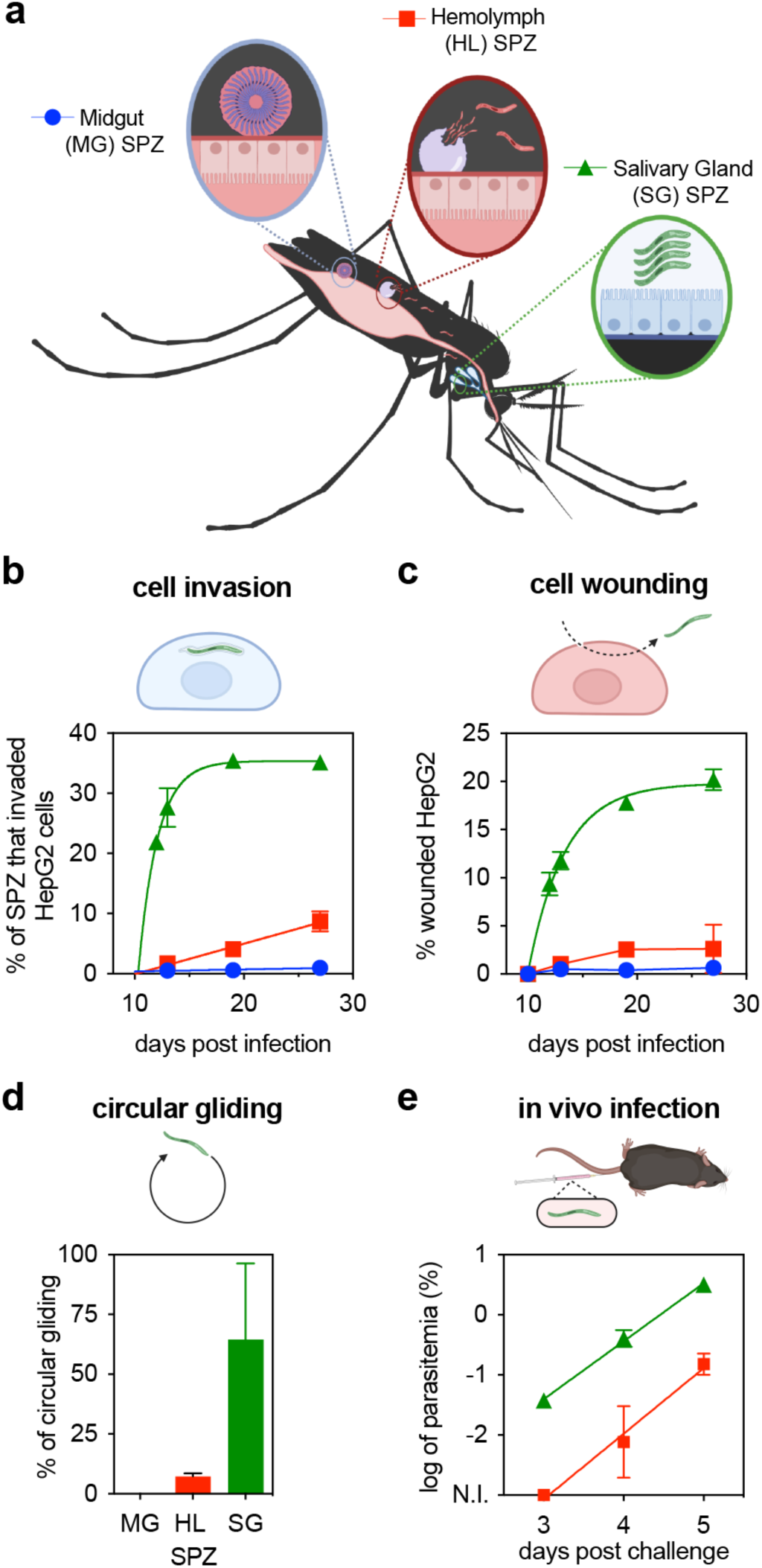
SPZ, formed in the MG, acquire maximal infectivity days after first invading the SG. **a.** Scheme showing the different forms of SPZ, forming inside oocysts on the mosquito MG (blue), egressing in the HL (red) and inside SG (green). **b-c.** Percentage of MG, HL and SG SPZ inside HepG2 cells (cell invasion, **b**) or percentage of wounded and resealed HepG2 cells (cell wounding, **c**) according to the day SPZ were collected across at least 3 mosquito infections. SG SPZ data followed a one phase decay non-linear fit in both cases (Supplementary Table 1). **d**. Infectivity differences can be explained by the reduction in gliding as measured by the percentage of MG, HL and SG SPZ isolated 14 days pi able to complete a full circle in 2D (Tukey’s multiple comparison test across at least 2 biological replicates). **e**. Comparison of the parasitaemia across time after intravenous injection of 5,000 HL or SG SPZ isolated 14 days pi in C57Bl/6 mice across 4 biological replicates. Linear regression of the log-transformed parasitaemia are represented. For all graphs, mean and standard deviation are represented.

As shown in Fig. 1b-c, MG SPZ exhibited minimal infectivity, with fewer than 0.25% successfully invading HepG2 cells. Consistently, the proportion of wounded HepG2 cells remained exceedingly low (<1.5%), corroborating the lack of expression of proteins involved in cell traversal in MG sporozoites (44,46,49). Upon egress into the HL, SPZ demonstrated a modest gain in invasive ability, reaching a maximum of 7.4% invasion by three weeks pi; however, traversal capacity remained poor, with wounded cells never exceeding 3% in average. By contrast, SG SPZ displayed a significantly greater capacity for both invasion and traversal, with 35% of parasites invading HepG2 cells and wounding 20% of cells, consistent with transcriptional reprogramming and acquisition of infectivity following SG invasion as described in prior studies (37,44–46).

Both anatomical location and time post infection influenced sporozoite functionality (Fig. 1b-c). HL and SG SPZ harvested at days 12 and 15 pi were significantly less competent in both invasion and traversal compared to those collected at day 18 pi or later. Logarithmic modelling of invasion and traversal rates for SG SPZ revealed a plateau phase around day 18 pi (R² = 0.99 and 0.94, respectively), indicative of a progressive acquisition of infectivity. Considering that in our experimental setup the number of SPZ within the SG rises until approximately 18 days pi before reaching a near steady state (Supplementary Fig. 1a), and that the proportion of invasive SPZ (∼35%) observed at day 18 pi approximates the number present at day 15 pi, it is likely that sporozoites require several days within the glands to achieve full maturation.

To complement these observations, we assessed SPZ motility, a function critical for both invasion and traversal (50,51). Motility assays performed at 14 days pi revealed that MG SPZ were essentially incapable of performing circular gliding under two-dimensional substrate, consistent with their poor infectivity. HL SPZ exhibited limited motility, with only 7.3% completing a full circle. In contrast, SG SPZ demonstrated robust, albeit variable, motility, with on average 65% of parasites executing at least one complete circle in two minutes of analysis (Fig. 1d).

Finally, to evaluate *in vivo* infectivity, we intravenously injected mice with 5,000 SPZ from day 14 pi, focusing on the two stages that have shown infectivity (HL and SG) and monitored subsequent parasitaemia. Consistent with *in vitro* observations, SG SPZ were highly infectious, with all mice displaying detectable parasitaemia by day 3 pi. By contrast, HL SPZ exhibited a one-day delay in patency, and by day 4 pi, parasitaemia in this group was approximately 1.5 logs lower than in mice infected with SG SPZ (Fig. 1e).

Together, these findings demonstrate that SPZ acquire full infectivity only after SG invasion, with MG SPZ remaining largely non-infectious. HL SPZ are in an intermediate functional state as they have acquired detectable infectivity, but remain markedly less competent than SG SPZ which progressively mature over several days, reaching optimal infectivity around 18 days pi.

### Sporozoites’ transcriptional maturation marker

*Uis4*, the most abundant transcript in *P.berghei* SG SPZ, displays increasing expression along pseudotime trajectories in single-cell RNA-seq analyses (44,46). To dynamically monitor SPZ maturation, we used a fluorescent reporter parasite line, which expresses mCherry under the control of the *uis4* promoter (52). This line allows measurement of *uis4* promoter activity via mCherry fluorescence, while expressing GFP under the control of the constitutive *eef1a* promoter, facilitating detection and flow cytometric analysis of SPZ (Fig. 2a).

**Figure 2.**
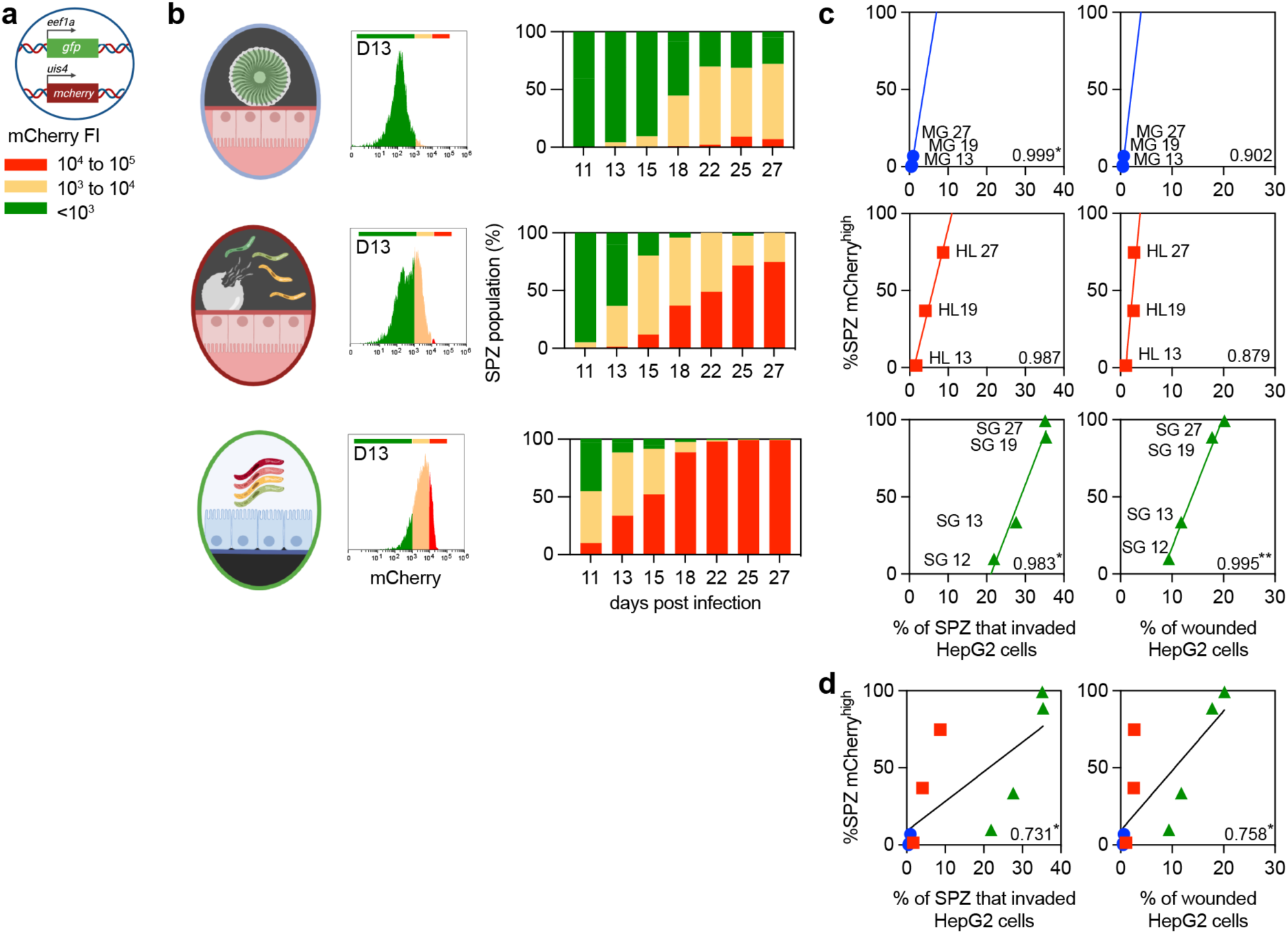
Measurement of SPZ maturity in tissues over time using a stage-specific fluorescent reporter. **a.** Fluorescence analysis of transgenic SPZs expressing GFP under the control of a constitutive promoter (*eef1a*) and mCherry under the control of a stage-specific promoter (*uis4*) were analysed by flow cytometry. Fluorescent intensity (FI) of the reporter was split into three categories: mCherry-high (10⁴-10⁵ arbitrary units (a.u.), red), mCherry-low (10³-10⁴ a.u., yellow), and mCherry-negative (<10³ a.u., green). **b.** SPZs were collected from MG, HL and SG in different days after the mosquito infectious feeding and cytometric analysis shows tissue and time-dependent increase of *uis4*-mcherry fluorescence from at least two independent mosquito transmissions. **c-d**. Percentage of SPZ expressing high levels of mCherry was correlated with the percentage of SPZ that invaded HepG2 cells (left) or the percentage of wounded cells (right) for each compartment (**c**) or altogether (**d**) by calculating their Pearson coefficients and significance (shown on the bottom right of each graph, * p < 0.05,** p < 0.01).

Using this line, we isolated MG, HL, and SG SPZ from the same infected mosquitoes and quantified GFP and mCherry fluorescence by flow cytometry from 11 to 27 days pi. GFP fluorescence remained stable across all stages, enabling consistent gating of SPZ based on GFP intensity and side scatter properties (Supplementary Fig. 1b). On the other hand, transcriptional activity measured by mCherry fluorescence in mCherry@*uis4* parasites increased as SPZ progressed from MG to HL and SG forms in the same mosquito as well as in each compartment over time (Fig. 2).

We further characterised the reporter line across the course of infection, stratifying mCherry fluorescence into three categories of intensity: mCherry^high^ (10⁴-10⁵, red), mCherry^low^ (10³-10⁴, yellow), and mCherry^neg^ (<10³, green). MG SPZ consistently failed to reach high levels of mCherry, whereas HL and, more prominently, SG SPZ showed progressive increase in mCherry expression over time. This observation reinforces the major transcriptional distinction between MG SPZ and the HL/SG SPZ populations, as highlighted in previous single-cell studies, and highlights a slow but steady and time-dependent transcription activation or de-repression (44,45).

Given the apparent temporal trend, we examined the relationship between the proportion of mCherry^high^ SPZ and their capacity to invade HepG2 cells. As shown in Fig. 2b, despite the small number of points, significant correlations were observed when analysing MG and SG SPZ individually (Fig. 2c, left panels, r > 0.98, p < 0.05* for MG and SG SPZ). However, when stages were pooled, the overall correlation weakened (Figure 2d, r = 0.731; p = 0.02*). Moreover, mCherry expression in SG SPZ at day 13 pi was comparable to HL SPZ at day 18-19 pi, despite displaying ∼7 times more invasion (Fig. 1b-c; Fig. 2c, left panels). Similar results were observed regarding cell traversal, except that only SG SPZ showed significant correlation between cell wounding and mCherry expression (r = 0.995, p = 0.01**; Fig. 2c, right panels) corroborating that only SG SPZ express the main proteins involved in cell wounding.

The linear regressions analysis also shows that MG and HL SPZ share a similar profile of mCherry/infectivity, with MG SPZ being proportionally less mature and infectious than HL SPZ. Acquisition of optimal motility, cell traversal and cell invasion activities is only observed after invasion of SG.

This finding highlights fundamental functional differences between different SPZ forms, even if they express similar levels of *uis4*-driven mCherry. Therefore, time-dependent *uis4* transcriptional maturation occurs in each mosquito compartment, but the maximal estimated infectivity of MG and HL SPZ (100% of mCherry^high^, <10% of invaders and <5% of wounded cells) is much inferior to the minimal infectivity of relatively immature SG SPZ at day 12 pi (10% of mCherry^high^, >20% of invaders and ∼10% of wounded cells).

Together, these data suggest that while *uis4* promoter activity serve as a useful marker of sporozoite maturation within each SPZ form, it does not reliably predict infectivity across different SPZ populations. In particular, late-stage HL SPZ can express high levels of *uis4-driven* mCherry yet remain markedly much less infectious than relatively immature SG SPZ.

### Salivated sporozoites display enhanced infectivity during early mosquito infection

We next turned our attention to salivated (SAL) SPZ. Although SG SPZ are routinely used in experimental infections, only SAL SPZ have the physiological opportunity to initiate infection in a mammalian host. Furthermore, single-cell transcriptomic analyses have revealed distinct transcriptional profiles between SG and SAL SPZ in *P. berghei* (44), which may help explain why mosquito bite-mediated infections are more efficient than direct intradermal injections of SG SPZ. Indeed, a small number of mosquito bites—delivering only a few tens to hundreds of SPZ—can establish blood-stage infections comparable to those resulting from intradermal injection of 5,000 SG SPZ (53).

To investigate this phenomenon, we used the mCherry@*uis4* line to compare mCherry fluorescence between SG and SAL SPZ. Saliva was collected by placing gel-loading tips containing 5 µL of 1% fatty acid-free BSA in DPBS around the mosquito proboscis and allowing salivation at 37°C for 15-20 minutes. As shown in Fig. 3a, SAL SPZ exhibited markedly higher mCherry fluorescence compared to SG SPZ 13 days pi, with no detectable mCherry^neg^ parasites, indicating elevated *uis4-driven* transcription. Control experiments confirmed that incubation at 37°C did not alter mCherry expression in SG SPZ (Supplementary Fig. 2a), suggesting that either mCherry^high^ SPZ are preferentially expelled or that the process of salivation itself induces transcriptional or translational changes.

**Figure 3.**
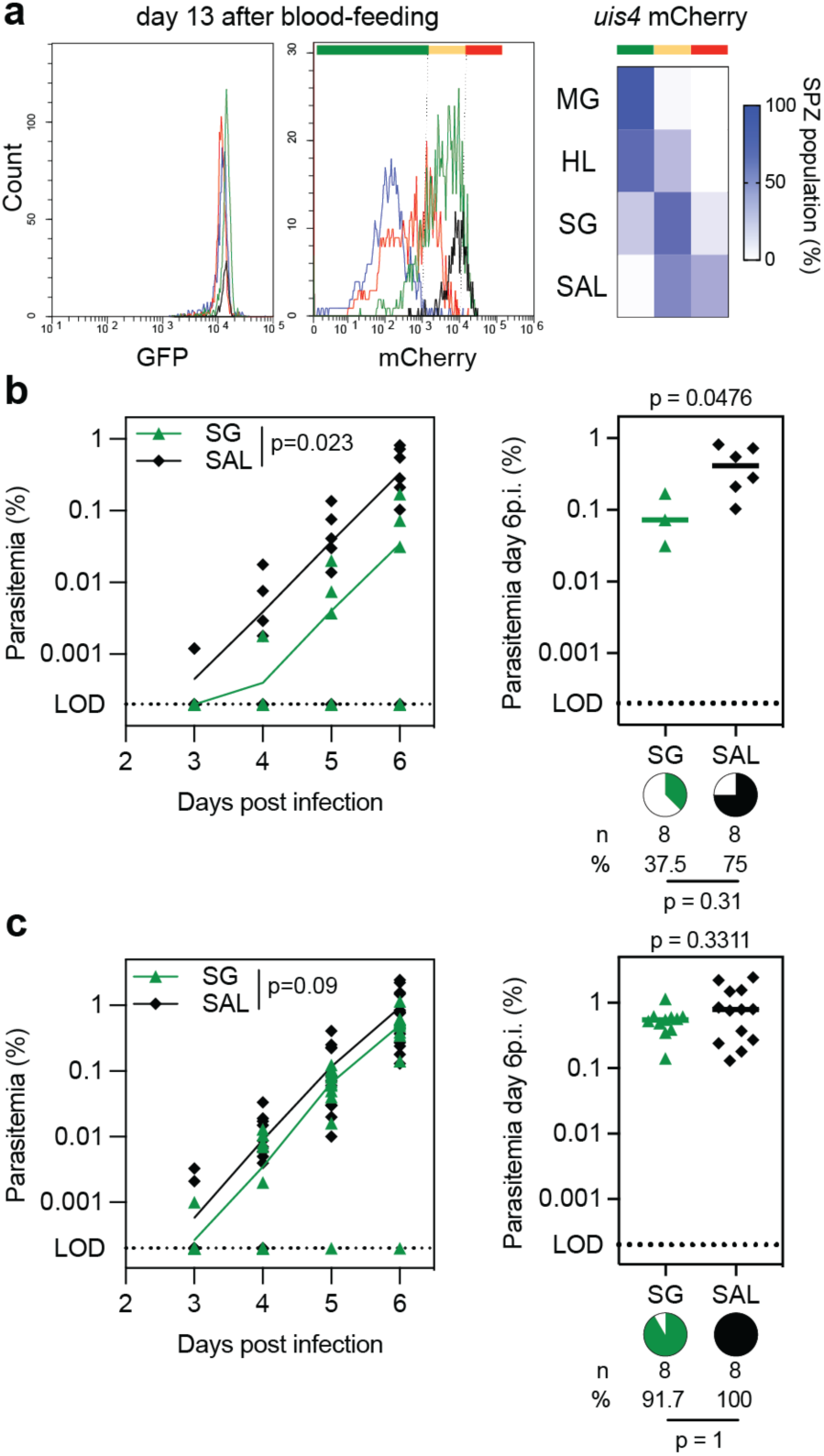
Salivated (SAL) SPZ are more infectious *in vivo* early in mosquito infection. **a**. Fluorescence analysis of MG, HL, SG and SAL SPZ (blue, red, green, and black respectively in histogram) collected 13 days post-infection (pi) summarised in a heatmap with percentage of SPZ expressing no (green; < 10^3^), low (yellow; between 10^3^ and 10^4^) or high (red; > 10^5^) levels of mCherry. **b-c**. 50 SG or SAL SPZ were isolated 15 days pi (**b**) or 250 SG or SAL SPZ were isolated 21 days pi (**c**) and injected IV in C57Bl/6 mice. Parasitemia was followed from 3 days pi (left graph) and parasitemia and prevalence of infection on day 6 pi were calculated (right graph). Median parasiteamia is represented in both cases and experiments were performed across 3 independent mosquito infections. Number of mice infected (n) and prevalence of infection (%, pie chart) are indicated for each sample. Parasiteamia across time were compared by two-way ANOVA (left graph), while parasiteamia and prevalence of infection on day 6 pi were compared by unpaired t-test and Fisher’s exact test respectively.

We next assessed whether the differences in transcriptional maturity correlated with infectivity *in vivo*. Infections were performed at 15 days pi and, due to the limited numbers of SAL SPZ available, only 50 SPZ were injected per C57BL/6 mouse intravenously. At this limiting inoculum, a proportion of mice remained uninfected, which reduces the precision of parasitaemia-based comparisons. Attempts to increase SPZ output by adapting feeder-based methods described in other studies proved unsuccessful as the isolated SAL SPZ were not infectious (Supplementary Fig. 2b). Parasitaemia post-IV injection was monitored daily from day 3 until mice reached 0.5% parasitaemia or day 10. As shown in Fig. 3b, mice infected with SAL SPZ exhibited significantly higher parasitaemia trajectories compared to those infected with SG SPZ (Two-way RM ANOVA, p = 0.02**). While prevalence of infection did not differ significantly—possibly due to the limited number of mice—parasitaemia at day 6 pi was significantly higher in SAL SPZ-infected animals (Mann-Whitney test, p = 0.048*).

We then repeated the comparative infections 21 days pi, when mCherry^high^ population is close to its maximum value in SG SPZ (Fig. 2b) and SG SPZ have reached their optimal infectivity (Fig. 1b-c). Increased SAL SPZ yields at this time point permitted the use of a higher inoculum (250 SPZ per mouse), selected to ensure infection in all mice and thereby enable quantitative assessment of subsequent blood-stage dynamics. As shown in Fig. 3c, although a trend towards greater infectivity of SAL SPZ remained, the differences did not reach statistical significance (Two-way RM ANOVA, p = 0.09).

### Salivated sporozoites form larger and more transcriptionally active liver stages *in vitro*

To assess whether the enhanced infectivity of SAL SPZ observed *in vivo* is reflected during liver-stage development and whether the trend observed after day 21 pi could be physiologically relevant, we performed *in vitro* infections of HepG2 cells using SAL and SG SPZ harvested at 19 days pi. Owing to this bottleneck, SAL and SG SPZ infections were performed at equivalent multiplicities of infection (MOI), with an additional SG control infection at a standard MOI of 1:4 (SPZ:HepG2 cell, high MOI). Quantification of EEFs at 48 hours pi revealed a ∼2.5 times more infected HepG2 cells in the SAL group relative to SG control (Fig. 4a, p = 0.045*), suggesting either increased efficiency of productive invasion or enhanced intracellular survival.

**Figure 4.**
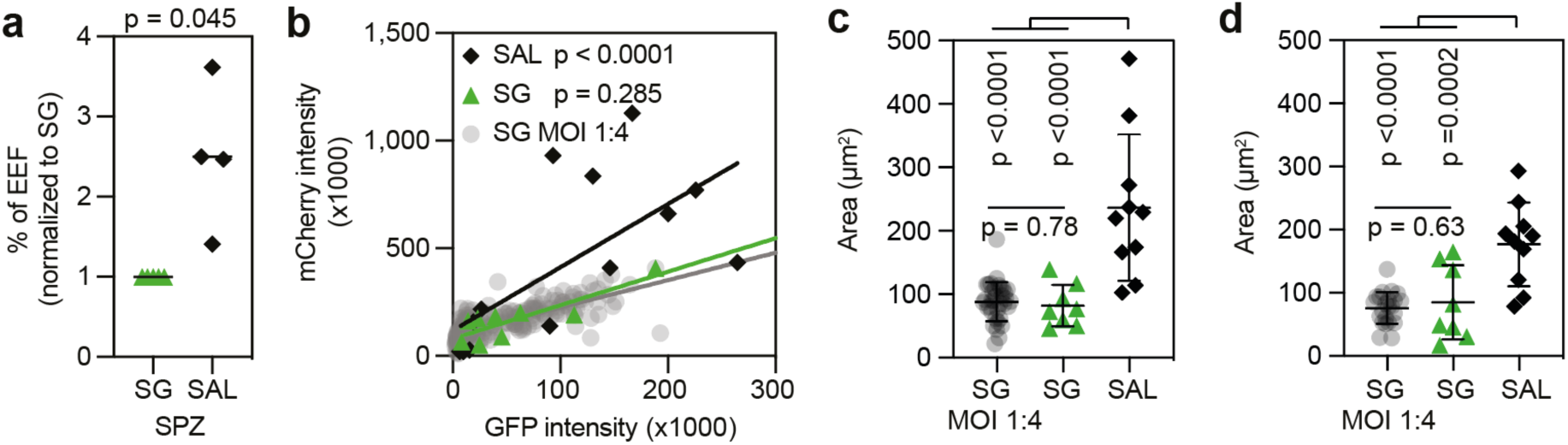
SAL SPZ are more efficient at infecting HepG2 *in vitro*, forming larger exo-erythrocytic forms (EEFs) **a**. Higher percentage of HepG2 cells are infected by SAL than SG SPZ when measured 48h pi. Four infections from independent transmissions were performed with 100 to 500 SPZ isolated 19 days pi depending of the rate of salivation (paired t-test). **b**. Comparison of linear regressions of GFP and mCherry fluorescence signals of 48h EEFs from SG and SAL SPZ using the same MOI, and from standard infection using SG SPZ at MOI 1:4 (SG MOI 1:4). P-values are from slope comparison with the SG MOI 1:4 regression using two Student t-tests and Bonferroni correction. **c-d**. Area of EEFs measured 48h pi using *mCherry@uis4* parasites (**c**) and GFP*@hsp70* parasites (**d**). Mean and standard error are displayed and adjusted P-values were obtained using Holm-Šídák’s multiple comparisons test.

To further explore the *uis4-driven* maturity marker in EEFs, we analysed the relationship between constitutively expressed GFP (*eef1a*-driven) and mCherry (*uis4*-driven) fluorescence in individual EEFs. Quantitative fluorescence analysis by flow cytometry indicated that EEFs derived from SAL SPZ exhibited significantly higher mCherry/GFP intensity ratio than SG SPZ at low and high MOI. Accordingly, the slope of the curve of EEFs derived from SG SPZ high MOI significantly differed from that using SAL SPZ (p <0.0001, Fig. 4b), but was similar to the slope of the linear regression using SG SPZ low MOI (p = 0.285, Fig. 4b). This indicated that the higher transcription of *uis4-*driven mCherry in SAL SPZ in comparison to SG SPZ (Fig. 3A) translated into a higher expression of this maturity marker also in liver-stages. This increased fluorescence ratio could also be related to an increase in parasite biomass, possibly reflecting more advanced development and/or larger EEFs. Quantification of EEF area at 48 hpi indeed revealed significantly larger EEFs in the SAL SPZ-infected cultures (Fig. 4c). To check if the observed phenotype was specific to the maturity reporter parasites, we repeated the experiment using *hsp70-*driven GFP SPZ, obtaining the same result (Fig. 4d).

Altogether, these findings suggest that SAL SPZ give rise to a higher number of, and transcriptionally more active, liver-stage parasites *in vitro,* even after SG SPZ have reached full infectivity. The transcriptional variability observed across individual EEFs further underscores distinct developmental dynamics between SG- and SAL-derived infections, warranting deeper investigation.

## Discussion

Sporozoite maturation has long been a subject of study across *Plasmodium* species and *Anopheles* mosquitoes, giving rise to discrepancies in reported infectivity and data interpretation. Transcriptomics studies comparing different SPZ forms—MG, HL, SG and occasionally SAL— have recently been published to investigate the transcriptional differences of these developmental stages, though no functional analysis was performed and correlated to the transcriptomics profiles (37,38,44–46). One of the key contributions of this study lies in its longitudinal approach. By characterising infectivity of all forms of SPZ throughout mosquito infection, we demonstrated that both tissue and time are essential to *P. berghei* SPZ maturation, reconciling different studies suggesting one or the other factor as a main or only driver for maturation (33,34,54,55). While HL SPZ reached some degree of infectivity with time, they remained sub optimally infectious, with lower *in vitro* infectivity and a delayed prepatent period when injected intravenously into mice, sharing a similar profile with MG SPZ, which is quite distinct from that of optimally infectious SG SPZ. Interestingly, SG SPZ also matured with time, reaching their peak infectivity and plateauing 18 days pi. Correlating with this time-dependent maturation, *uis4-*driven transcripts level can be used as a marker for infectivity, though only within SPZ forms as similar *uis4* expression in HL and SG SPZ results in dissimilar motility, invasion and cell traversal activities. More broadly, reporter-based tracking of maturation state may provide a useful framework for optimising axenic sporozoite production systems, particularly in *P. falciparum*, where functional maturation remains difficult to assess directly. A previous extensive study on the HL infectivity and ability to trigger a protective immune response in the mouse (56) have shown similar phenotypes for this population. They found HL SPZ to be mostly non motile in 2D (about 14% vs. 7.5% in the current study), with limited cell wounding activity as they were not detectably above the no-sporozoite control. Regarding *in vivo* infection, they observed a ten-fold reduction in parasitaemia 4d post-intravenous injection of HL or SG SPZ, slightly below our 1.5 log difference. Differences in the *in vitro* invasion assay methodology, microscopy read out of Huh7 infection in their case and our flow cytometry analysis of HepG2 invasion, make any comparison unsuitable. The concordance of their phenotype results with ours, in addition to the different flow cytometry profiles of MG and HL (Fig 2c and Supplementary Fig. S1b) indicate the robustness of the HL SPZ isolation method and the limited, if any, contamination by MG SPZ.

Finally, SAL SPZ showed enhanced infectivity compared to SG SPZ, suggesting that the salivated population may be enriched for parasites that have progressed further along the maturation programme. Given that salivary gland invasion can begin several days before our earliest salivation collections (2), SPZ recovered by salivation at early time points are likely biased towards “early invaders” that have resided longer within the glands, matured further, and reached the ductal region from which SPZ are preferentially salivated. Consistent with this, increased motility in the mature SG SPZ population has been associated with the colonization of the mosquito salivary ducts and could explain the preferential salivation of this more infectious population (12). An alternative explanation could be that salivation-associated cues could contribute to the maturation process, noting that these possibilities are not mutually exclusive. While the characterisation of this population was limited by the number of SPZ that could be obtained, the preliminary data provided here indicate that SAL SPZ are more likely to establish EEFs and that the resulting liver stages are larger and more transcriptionally active. The precise magnitude and generalisability of these SAL-associated phenotypes will, however, require further study across additional experimental settings and larger datasets. The increase in EEF numbers is consistent with elevated *uis4* transcript level in SAL SPZ and with a role for UIS4 in evading host clearance, suggesting that reduced elimination may contribute to the higher frequency of productive liver-stage infections (57). The increased size and transcriptional activity of SAL-derived EEFs further suggest a kinetic advantage after hepatocyte entry. In this context, recent work reporting that salivated SPZ contain proportionally less mRNA but more rRNA (44) raises the possibility that SAL SPZ possess greater translational capacity at baseline, which could support more rapid establishment of liver-stage growth and ultimately manifest as larger, more transcriptionally active EEFs. Whether other mechanisms—such as enhanced invasion or egress of more liver-stage merozoites—contribute to higher parasitaemia in SAL-infected mice remains to be determined.

SG invasion remains a poorly understood mechanism, with few vector-parasite interaction identified (5,36,40,58). The key role of rhoptry, invasive organelles shown to be essential to hepatocyte and red blood cell invasion, has long been studied though only recently was confirmed as SG SPZ are found to have fewer rhoptries than MG and HL SPZ (8,59–63) and knockout or knockdown of rhoptry proteins impair SG invasion (8,60–63). This clear distinction between SG SPZ to the previous forms of the parasite is likely related to the transcriptional and translational changes observed in SG SPZ such as increased *uis4* transcripts levels, as well as the global changes described by others including the high expression of cell traversal proteins (37,38,44,46,64).

Interestingly, SG invasion is essential but not sufficient for maturation as *P. yoelii* SPZ that could invade *An. albimanus* SG could not establish a liver infection in mice suggesting that other environmental factors are at play than solely rhoptry discharge (65). Additionally, sporozoite maturation is time-dependent, as was first noted in *P. gallinaceum* (54,55), but appears to plateau with only ∼30 % of SG SPZ capable of infecting hepatocytes *in vitro* at any given time, suggesting that not all sporozoites attain full infectivity. Why only a subset of SG SPZ is competent for invasion, and whether this correlates with their spatial positioning within the gland, remains to be determined. Recent advances in expansion microscopy and spatial transcriptomics, combined with our newly characterised fluorescent reporter line, offer promising tools to begin resolving the anatomical and molecular determinants of sporozoite infectivity at single-parasite resolution (59,66).

Notably, our data differ from a similar study looking at P*. falciparum* SG SPZ infectivity (67). While we found no decrease in *P. berghei in vitro* infectivity from 18 days pi to 28 days pi, van Schuijlenburg and colleagues found that *P. falciparum* SPZ slowly “age” and become less motile, infectious and immunogenic. Single-cell transcriptomic analyses support this distinction: while both the Bogale and Real datasets describe stage-dependent transcriptional reprogramming, the timing and nature of these shifts differ between species. *P. berghei* SPZ appear to maintain a quiescent state within the SG more effectively and are able to activate rapidly upon salivation. This is evident both in scRNA-seq data—reflected in a higher proportion of ribosomal RNA and markers of translational activation—and in our functional assays (44). On the other hand, Real et al. have found little to no difference in transcription comparing SG SPZ and SAL SPZ (45). These biological differences are echoed in practical handling: while *P. berghei* SG SPZ are typically dissected in PBS with dissection lasting for several hours, *P. falciparum* require maintenance in insect medium and lose infectivity within 1-2 hours. Together, these findings suggest that *P. berghei* sporozoites are primed for rapid activation and accelerated protein synthesis upon salivation, a feature that may facilitate efficient nutrient acquisition and successful transition to liver-stage development.

Further optimisation of the salivation protocol will be critical to enable a more detailed and robust comparison between SG and SAL SPZ and to elucidate how their functional and molecular differences influence liver-stage infection. When mosquitoes were allowed to salivate into feeders, no infectivity was observed—possibly due to excessive dilution, reducing saliva proteins concentration and SPZ-SPZ proximity cues that may help preserve viability. Enhancing SPZ yield and concentration would not only facilitate more robust intravenous infections but also enable intradermal injections, which are more physiologically relevant. Such a protocol would likely exacerbate any phenotype linked to salivation, especially in later timepoints where the functional divergence between SAL and SG SPZ may become more subtle. Beyond infection rates, further characterisation will be needed to determine whether the observed increase in parasitaemia following SAL SPZ injection reflects greater hepatocyte invasion, reduced clearance, or both. Analysis of merosome output and quantification of liver-stage merozoite release could provide mechanistic insight into the basis of this *in vivo* phenotype. An additional variable likely to merit attention is the mosquito background itself. Differences between laboratory colonies, or even between mosquito species, may influence the kinetics and extent of SPZ maturation, given the long-term independent maintenance and divergence of many experimental populations. A particularly attractive avenue for future work would be a comparative analysis across mosquito populations, species, or even individual mosquitoes, to identify vector backgrounds that favour accelerated SPZ maturation or the generation of more highly infectious parasites.

Ultimately, understanding how sporozoites acquire infectivity—and the environmental and molecular triggers that orchestrate this transition—is essential for the rational development of axenic SPZ. Defining the sequence of events required for functional maturation will not only illuminate fundamental aspects of *Plasmodium* biology but also support efforts to produce transmission-competent SPZ for vaccine and therapeutic applications in the absence of a mosquito vector.

## Method

### Ethics statement

Experiments involving animals were all approved by the Animal Care and Use Committee of Institut Pasteur (CETEA Institut Pasteur 2013-0093, Ministère de l’Enseignement Supérieur et de la Recherche MESR 01324) and performed according to European guidelines and regulations (directive 2010/63/EU). Mice were sourced from Janvier Labs.

### Parasites and mosquito infections

Different reporter parasite lines were used in this study, all generated in the *P. berghei* ANKA background. A strong fluorescent line that expresses GFP under the *hsp70* promoter (68) was used for *in vivo* experiments to facilitate the measure of blood parasitaemia. The other reporter line used was a *P. berghei* ANKA expressing GFP under the *eef1a* promoter background: a transcription reporter generated by introducing mCherry driven by *uis4* 5’ and 3’ UTR (52).

All mosquitoes used were *Anopheles stephensi* mosquitoes (SDA500 strain) that were reared in the Centre for Production and Infection of Anopheles (CEPIA) at Institut Pasteur and maintained on 10% sucrose/water. For production of *P. berghei* SPZ, 4-week old RjOrl:SWISS were infected by IP injection of infected red blood cells. Parasiteamia was left to establish over 3-4 days until mature gametocytes were present and mice were used to infect 1 to 2-day old mosquitoes. One week post infectious blood meal, mosquitoes were fed on naïve 5-week old RjOrl:SWISS mice.

### SPZ isolation and flow cytometry analysis

SPZ isolation was performed in DPBS 1X as early as 11 days post infectious bloodmeal, and up to 28 days pi. MG SPZ were obtained by crushing infected MG with a pestle, releasing SPZ from the oocysts. HL SPZ were isolated by cutting the last segments of a mosquito’s abdomen and washing the HL out by gently flushing 10-20 µL of DPBS with an insulin syringe introduced into the mosquito thorax. SG SPZ were isolated from infected glands and freed after crushing them with a pestle. In all cases, mosquito debris were removed by filtrating through a 35 µm filter before viable GFP^+^ sporozoites were counted by fluorescent microscopy or flow cytometry. When analysing the fluorescence by flow cytometry, sporozoites were simply diluted to less than 10,000 SPZ per microliter

### Mosquito salivation

Mosquitoes were aspirated, knocked down on ice and sorted under a fluorescent stereomicroscope to select insects with infected SG. They were then attached on a microscope slide by gluing their wings to tape. Their proboscises were encased into a gel loading tip containing 5µL of 1% fatty acid-free BSA/DPBS (Sigma-Aldrich A6003). Mosquitoes were left to salivate 20 min at 37°C in a humidified chamber and tips’ content was pooled before counting GFP^+^ sporozoites on Kova under fluorescence or by flow cytometry using CountBright™ Absolute Counting Beads (Invitrogen C36950). To account for different handling, all SPZ used in head-to-head comparison to SAL SPZ were dissected in 1% fatty acid-free BSA/DPBS, diluted to the same concentration as the SAL SPZ, and incubated at 37°C for 20 min before proceeding.

### Sporozoite motility assay

All materials were kept on ice until they were placed in the microscope chamber. Freshly isolated SPZ collected 14 days pi were analysed immediately after isolation. GFP^+^ sporozoites were counted, resuspended in DPBS 1X containing 10% heat inactivated foetal bovine serum (FBS) and 5,000 SPZ were seeded in an 18 well ibiTreat μ-slide well. Parasites were gently centrifuged on the slide at 500 x *g* for 3 min at 4°C. The slide was then transferred to an inverted Axio Observer Z.1 microscope (Zeiss) chamber prewarmed at 37°C and under 5% CO_2_. Temperature was left to equilibrate for 5 min before short live fluorescence recordings were obtained with an image taken every second for 2 min. Motility was scored manually from the recordings on Fiji (69), and sporozoites were classified as motile only when they completed at least one full circular trajectory during the observation period. Motility was expressed as the percentage of GFP^+^ sporozoites that completed at least one full circle among the total number of GFP^+^ sporozoites observed in each preparation.

#### Cell maintenance and *in vitro* assays set up

HepG2 cells (ATCC HB-8065) were maintained on DMEM high glucose containing GlutaMAX (Gibco 10566016) completed with 10% FBS, 1X of non-essential amino acids, 2% penicillin-streptomycin-neomycin. Cells were passaged when reaching confluency or every 3-4 days. *In vitro* assays were performed as described before (70). Briefly, 40,000 HepG2 cells were seeded per well in 96-well plates a day prior to infection. SPZ were isolated as described above. After filtration to remove debris and count of viable GFP^+^ SPZ, they were resuspended in complete DMEM high glucose media and added to the cells at an MOI of 1:4 (unless specified otherwise). In the case of cell wounding assay, dextran-rhodamine (Invitrogen, D1817) was included to a final concentration of 1 mg/mL. Parasites were then gently centrifugated onto cells at 100 x g for 3 min and incubated at 37°C 37 °C, 5% CO_2_ and 10% O_2_.

### Cell wounding and invasion assay analysis

At 2h pi, culture supernatants were collected and cells were washed in DPBS, collected as well. Cells were trypsinised and pooled with the recovered media before analysis by flow cytometry (CytoFLEX S, Beckman). Extracellular sporozoites recovered from the wells were identify as GFP^+^ events with low side scatter outside of any gate. HepG2 single cells were then identified on the basis of forward and side scatter characteristics. Wounded cells were then quantified as percentage of dextran-rhodamine^+^ events within the HepG2 gate and invaded cells were identified as GFP^+^ events within the HepG2 population. Cell invasion, or the percentage of SPZ that have invaded, was calculated as the number of intracellular GFP^+^ parasites divided by the total number of GFP^+^ parasites recovered, calculated as the sum of intracellular GFP^+^ HepG2-associated parasites plus extracellular GFP^+^ sporozoites.

### EEF development and analysis

For EEF assays, cells were washed 2 h post-infection to remove non-invaded parasites and then cultured for a further 46 h with daily medium replacement. At 48 h post-infection, infected cells were first imaged with an inverted Axio Observer Z.1 microscope (Zeiss) followed by microscopy analysis on Fiji (69). Individual fluorescent liver stages were identified by their GFP fluorescence, and parasite area was measured for each EEF.

The cells were then washed and detached with trypsin, for flow cytometry analysis (CytoFLEX S, Beckman). EEF burden was quantified as the percentage of GFP^+^ HepG2 cells. In experiments performed with the mCherry@*uis4* reporter line, GFP fluorescence driven by the constitutive *eef1a* promoter and mCherry fluorescence driven by the *uis4* regulatory regions were recorded for each infected cell, allowing comparison of the GFP/mCherry fluorescence relationship between conditions.

### SPZ challenge

As described previously (70), four- to five-week-old C57Bl/6 mice were used for *in vivo*infections. SPZ were collected as described above, purified by filtration and live parasites were counted using their fluorescence. Unless specified otherwise, 1,000 SPZ were injected intravenously in the mice tail vein. Parasitaemia was followed from 3 days to 10 days pi, or until parasiteamia reached 0.5%, by collecting a drop of blood from the tip of the tail in PBS and analysing samples by flow cytometry (CytoFLEX S, Beckman). GFP fluorescence was used to identified parasitised red blood cells and 250,000 to 500,000 erythrocytes were recorded.

## Acknowledgements

The authors would like to thank the Center for Production and Infection of Anopheles (CEPIA, C2RA, Institut Pasteur) for providing all mosquitoes used in the study and the Central Animal Facility (C2RA, Institut Pasteur) for caring for the mice.

---

**Supplementary Figure 1.**
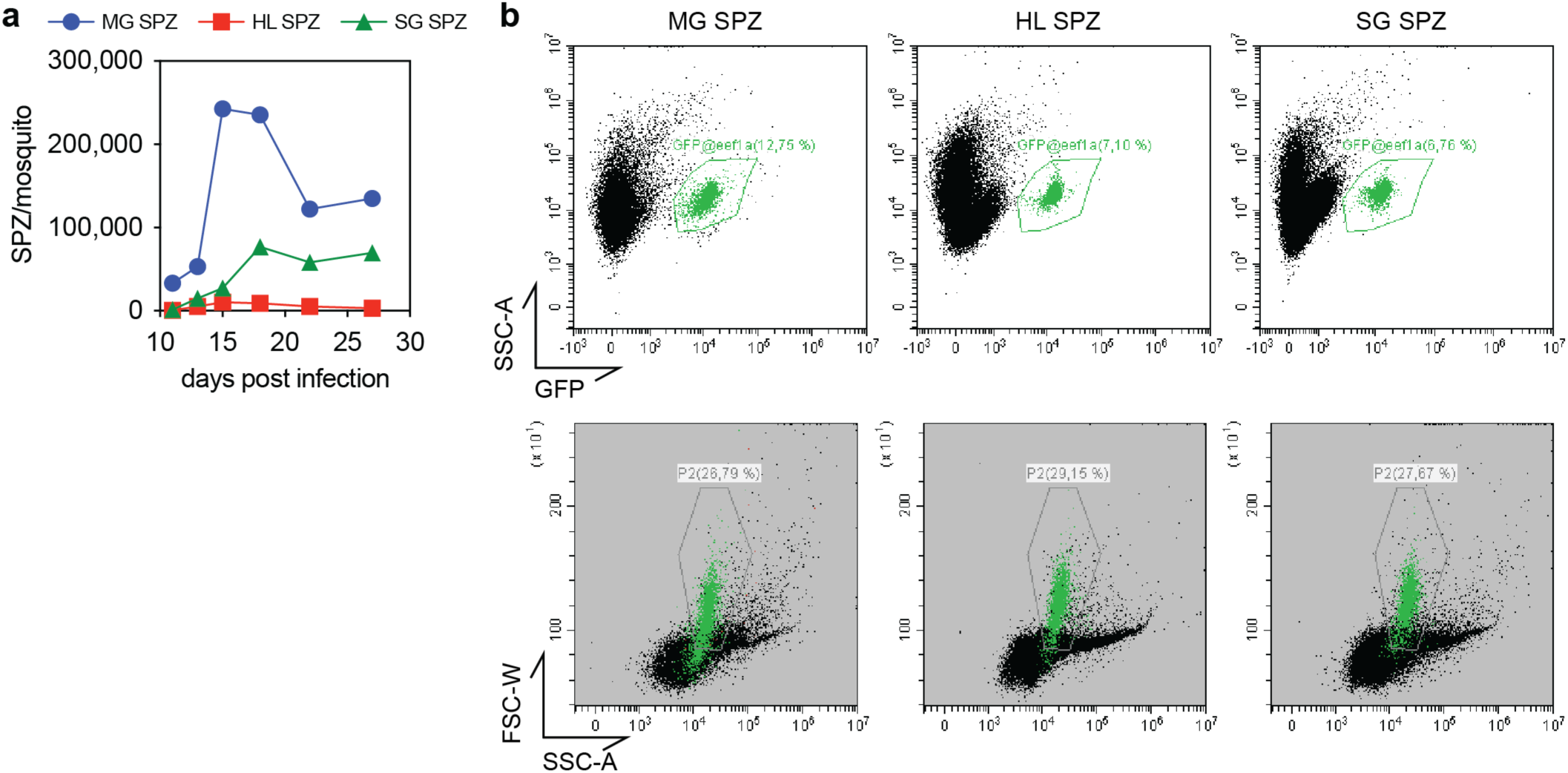
Dynamics of the different SPZ forms number and morphology. **a**. Number of SPZ from each compartment throughout mosquito infection. **B.** SPZ expressing GFP constitutively under the *eef1a* promoter were gated according to their green fluorescent; the resulting profile by side scatter area (x axis, logged) and forward scatter width (yaxis) is displayed in green and show a reduction in the spread of forward scatter values, with SG and SPZ having higher values.

**Supplementary Figure 2.**
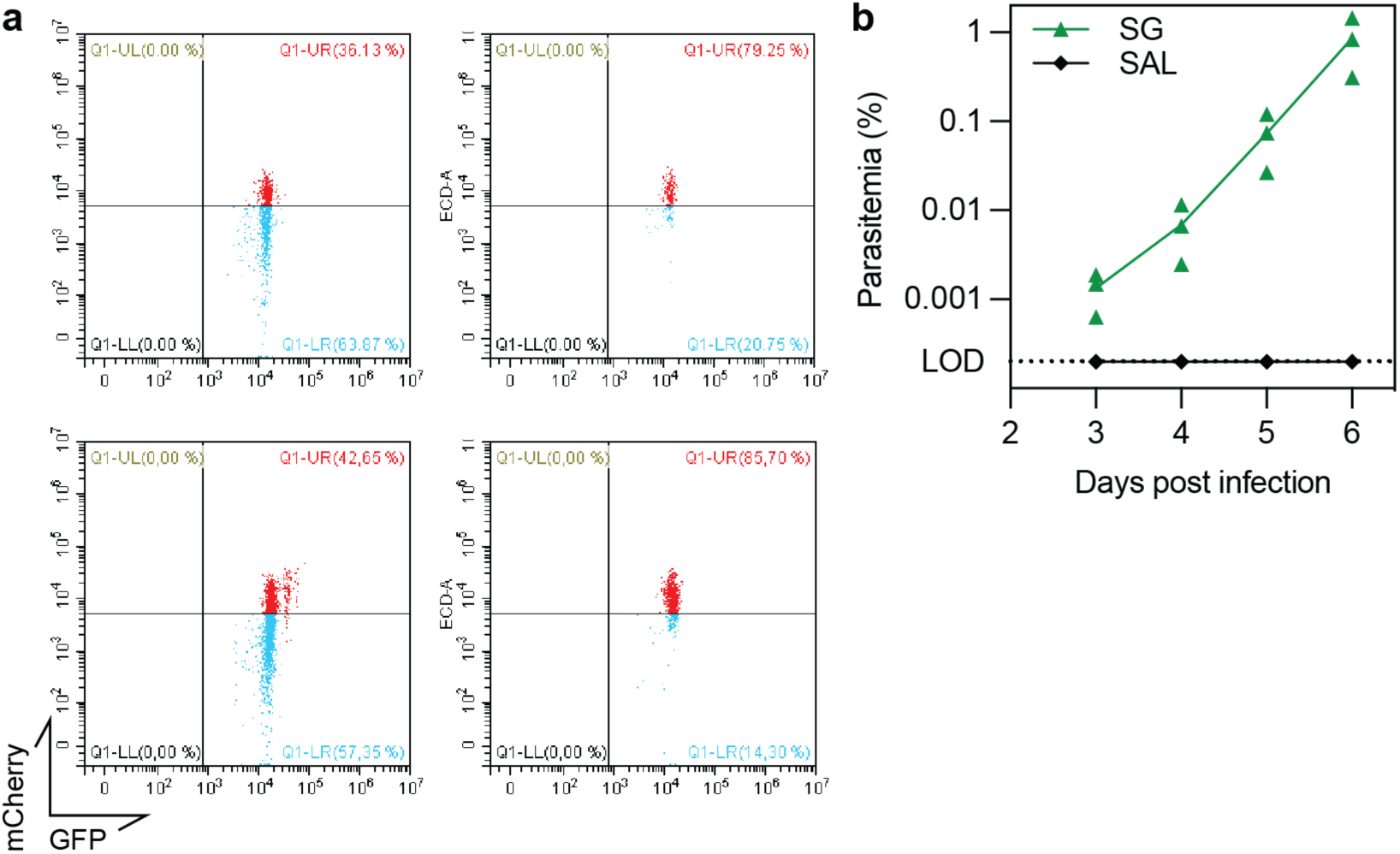
Temperature does not induce a shift in *uis4* transcription while salivation protocol can lead to loss of infectivity. **a**. Temperature does not induce a shift in uis4 transcription in SG and SAL mCherry@*uis4* SPZ were collected and gated as show in Supplementary Fig. 1a. Difference mCherry fluorescence profiles were observed (top) and which were unchanged after incubation at 37°C for 20 min, condition of salivation (bottom), confirming the higher mCherry expression in SAL SPZ was not due to the higher temperature they were exposed to. **b**. Salivation protocol can lead to loss of infectivity. Infected mosquitoes were left to feed on 1% fatty acid-free BSA/DPBS for 10 min before count and injection into 3 four-week-old C57Bl/6 mice. No mice developed a blood stage infection when injected with SAL SG while control mice who received SG SPZ were all positive 3 days pi.

**Supplementary Table 1.** Statistical parameters of the one phase decay non-linear fit of SPZ infection across time and compartment.

|  | Cell invasion |  |  | Cell traversal |  |  |
| --- | --- | --- | --- | --- | --- | --- |
|  | MG | HL | SG | MG | HL | SG |
| **Best-fit values** |  |  |  |  |  |  |
| Y0 | 0,05810 | -5,133 | -12341 | -2301 | -18,70 | -373,4 |
| Plateau | Unstable | Unstable | 35,39 | 0,5272 | 2,833 | 19,81 |
| K | 0,0001990 | 5,616e-005 | 0,5681 | 0,8382 | 0,1972 | 0,2995 |
| Half Life | 3484 | 12342 | 1,220 | 0,8269 | 3,515 | 2,314 |
| Tau | 5026 | 17806 | 1,760 | 1,193 | 5,070 | 3,339 |
| Span | Unstable | Unstable | -12377 | -2301 | -21,53 | -393,2 |
| R squared | 0,4604 | 0,9100 | 0,9355 | 0,2113 | 0,3872 | 0,9874 |

